# Amitosis as a strategy of cell division - Insight from the proliferation of Tetrahymena thermophila macronucleus

**DOI:** 10.1101/2020.08.11.247031

**Authors:** Y.X. Fu, G.Y. Wang, K. Chen, X.F. Ma, S.Q. Liu, W. Miao

## Abstract

Cell division is a necessity of life which can be either mitotic or amitotic. While both are fundamental, amitosis is sometimes considered a relic of little importance in biology. Nevertheless, eukaryotes often have polyploid cells, including cancer cells, which may divide amitotically. To understand how amitosis ensures the completion of cell division, we turn to the macronuclei of ciliates. The grand scheme governing the proliferation of the macronuclei of ciliate cells, which involves chromosomal replication and the amitosis, is currently unknown. Using a novel model that encompasses a wide range of mechanisms together with experimental data of the composition of mating types at different stages derived from a single karyonide of *Tetrahymena thermophila*, we show that the chromosomal replication of the macronucleus has a strong head-start effect, with only about five copies of chromosomes replicated at a time and persistent reuse of the chromosomes involved in the early replication. Furthermore the fission of a fully grown macronucleus is non-random, with a strong tendency to push chromosomes and their replications to the same daughter cell. Similar strategies may exist for other Tetrahymena species or ciliates, and have implications to the amitosis of polyploid cells of multicellular organisms.

---

Living organisms need cell divisions to grow or just to stay alive, which can be accomplished in two ways: mitosis and amitosis. The former is characterized by the presence of a spindle structure which pulls apart original and duplicated chromosomes in a cell to ensures the fidelity of daughter cells to its parental one, and the latter characterized by devoid of a spindle structure, therefore identity of the two daughter cells is less certain. It is well known that prokaryotic cells divide amitotically (binary fission) while most eukaryotic cells divide mitotically. Although both mechanisms are fundamental the amitosis is often regarded as a relic of little importance in biology. However, recent studies suggest that amitosis may be considerably underappreciated for multi-cellular eukaryotes^1,2^.

For eukaryotes, amitosis occurs predominantly in polyploid cells (that is, containing 3 or more copies of its haploid chromosome set), which are common throughout nature, ranging from bateria^3^, single cell eukaryote ciliates^4^, plants^5,6^ and also in some tissues of diploid animals including humans^7^. Drosophila, one of the most studied multicellular organisms for ploidy, has cells in salivary gland with ploidy level as high as 20489 while mouse giant trophoblast cells may have ploidy level up to 859^7^. For normally diploid organisms, polyploid cells are known to be generated by endoreplication, which is a variation of the mitosis process resulting in the increase of chromosomal numbers without cell division^8–10^. Much focus has been paid to the molecular mechanism that leads to endoreplication and the function of such polyploid cells, including tissue regrowth^7,11–13^, disease resistance/causation (such as in cancer)^2^ and regeneration of tissue specific stem cells^1^. Polyploid plant cells usually can divide mitotically, while the division of animal polyploid cells is more complex. They may lead to polyploid^14,15^ or depolyploidation resulting in diploid daughter cells. Although the division of polyploid cells is in general not well characterized, amitosis is likely an important mechanism for normal development to many eukaryotes judging from the observations in ciliates^16^ and Drosophila intestine^1,2,17^, and for cancer-causing in mouse^18^. However, amitosis leads to uncertainty of the genetic makeup of daughter cells, which opens up many possibilities that are poorly understood. Furthermore, whether the growth of a high ploidy cell is a simple process of duplicating all its chromosomes equally is unknown, which clearly can not be true for a cell of ploidy level that is not equal to a multiple of its haploid chromosome number. Therefore, the proliferation of polyploid cells is a fundamental and common process across nature with many mysteries remaining.

The single cell eukaryote ciliates are important model organisms^19–21^, characterized by the presence of two types of nuclei, one small diploid micronucleus and one large polyploid macronucleus^4,22–24^. The former serves as the germline nucleus and is not active during asexual reproduction, while the latter originates from the micronucleus during sexual reproduction, and goes through heavy editing and amplification to become high ploidy. Unlike most eukaryotes, the macronuclei of ciliates divide amitotically. While the propagation of the micronucleus follows the same pattern of typical eukaryotic cell mitosis, that is, each chromosome is doubled and one is pulled to each end of the nucleus and a split results in two identical daughter nuclei, much of the molecular mechanism governing the chromosomal replication and fission of the macronucleus remains unknown^25^. Nevertheless, ciliates provide an excellent model for studying the proliferation of polyploid cells. The answers to the puzzle is not only fundamental for ciliates, but have important ramifications to the polypoid cells in multi-cellular organisms, including humans. To date, the fission is assumed to be at random^26–30^ which was partly derived with the assumption that each chromosome in the macronucleus is duplicated once. Although random fission appears to be compatible with some historical data^28^, its profound consequence of requiring a lengthy process for purifying the mating type in a cell is overwhelmingly incompatible with observations. The inadequate understanding of the amitosis has become a serious obstacle for developing a proper population genetics model for analyzing DNA polymorphism in Tetrahymena.

Although the grand scheme of the proliferation of ciliate macronuclei is largely unknown after decades of research, the presence of multiple mating types (mts)^30–33^, coupled with recent molecular characterization^34–37^ in the most well-known ciliate, *Tetrahymena thermophila* (*T. thermophila* hereafter), provides intriguing opportunities to understand the mechanism through the tracking of mating type determination (mtd) chromosomes. There are seven mating types in *T. thermophila* and a karyonide which has a fresh macronucleus generated from the micronucleus has the potential to lead to a clonal population of single or multiple mts depending on the composition of its initial mtd chromosomes. A karyonide will go through many generations (divisions) to become sexually mature and at maturity, a cell is generally thought to contain a single mating type(mt) chromosome in its macronucleus. Decades of empirical observations show that when mature, the proportion of pure clones derived from a single karyonide is more than 50% or higher. A series of attempts have been made to explain the phenomenon which inevitably involved further specification and hypotheses about the binary fission process^30,38,39^. The latest incarnation of thought is that the phenomenon is in essence due to a combination of head-start replications with intranuclear coordination^30^. The head-start mechanism assumes that during the growing phase of proliferation, some chromosomes have acquired an advantage over the others early on so they are replicated multiple times with the expense of some of those not replicating. However this as well as some alternative mechanisms have not been subjected to rigorous scrutiny.

The clonal population derived from a single karyonide at any time consists of cells with macronuclei that are the product of successive rounds of growth and fission. Therefore, the composition of the macronuclei is the product of the process when the karyonide contains multiple mts. Here we propose a novel model of proliferation which is specified by three parameters. We conducted a series of experiments to track the compositions of mts of a clonal population derived from a single karyonide of *T. thermophila*, then the inference about the models was performed through a likelihood framework which led to new insights into the strategy of proliferation.

## The model for macronucleus proliferation

In the process of cell proliferation, the Tetrahymena macronucleus undergoes both chromosomal replications and fission. The former leads to the doubling of chromosomes within the macronucleus, the latter leads to segregation of the chromosomes in the fully grown macronucleus into two halves, each forms the macronucleus of a daughter cell. The process must have a molecular basis and the understanding of which will be of great importance, however, stochasticity is undoubtedly an important feature and its characterization, which we refer to as the grand strategy of proliferation, is at the center of modeling.

Assume that in the chromosomal replication phase, chromosome replications proceed in batches due to limited availability of concurrent replication apparatuses. The number of *concurrent replication apparatuses* likely differs among species and may also vary to some extent in different environments within the same species, but for simplicity it is assumed to be a constant *κ*. Next, each chromosome is associated with a *dominance level ω*. The probability of the *i*-th chromosome being selected into the batch for the next round of replication is equal to *ω*_*i*_/ ∑_*i*_*ω*_*i*_ where summation is taken over all eligible chromosomes. Initially all chromosomes have an equal dominance level 1, and after each round of replication, both the *κ* chromosomes in the current batch of replication and their *κ* replicates have their dominance levels changed to *ω*, while all other chromosomes have their dominance levels equal to 1. The replication continues until the desirable number of chromosomes is reached. In the fission phase, it is assumed that the fully grown set of chromosomes is initially split randomly into two halves, and the one with the most frequent (dominant) mt will increase by up to *ϕ*, which will be referred to as *shift*, chromosomes by exchanging non-dominant types with the dominant mt chromosomes in the other half. Once the exchange is completed, the macronucleus splits and each half forms the macronucleus of one daughter cell. It should be emphasized that the split process described above serves as a proxy for the underlying mechanism, and *ϕ* is a simple measure of the degree of departure from random split. The model is not intended to imply the actual mechanism of the cell split, which illumination requires future molecular experiments. However, it is imaginable that due to the physical proximity of templates and their duplicates, even a random cleavage of a cell may lead to a non-random split of the existing chromosomes. Steps in the model are illustrated in Figure 1. Throughout the paper, we shall refer to the collection of the three parameters of the model as ***θ***. That is

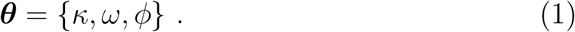

**Figure 1.**
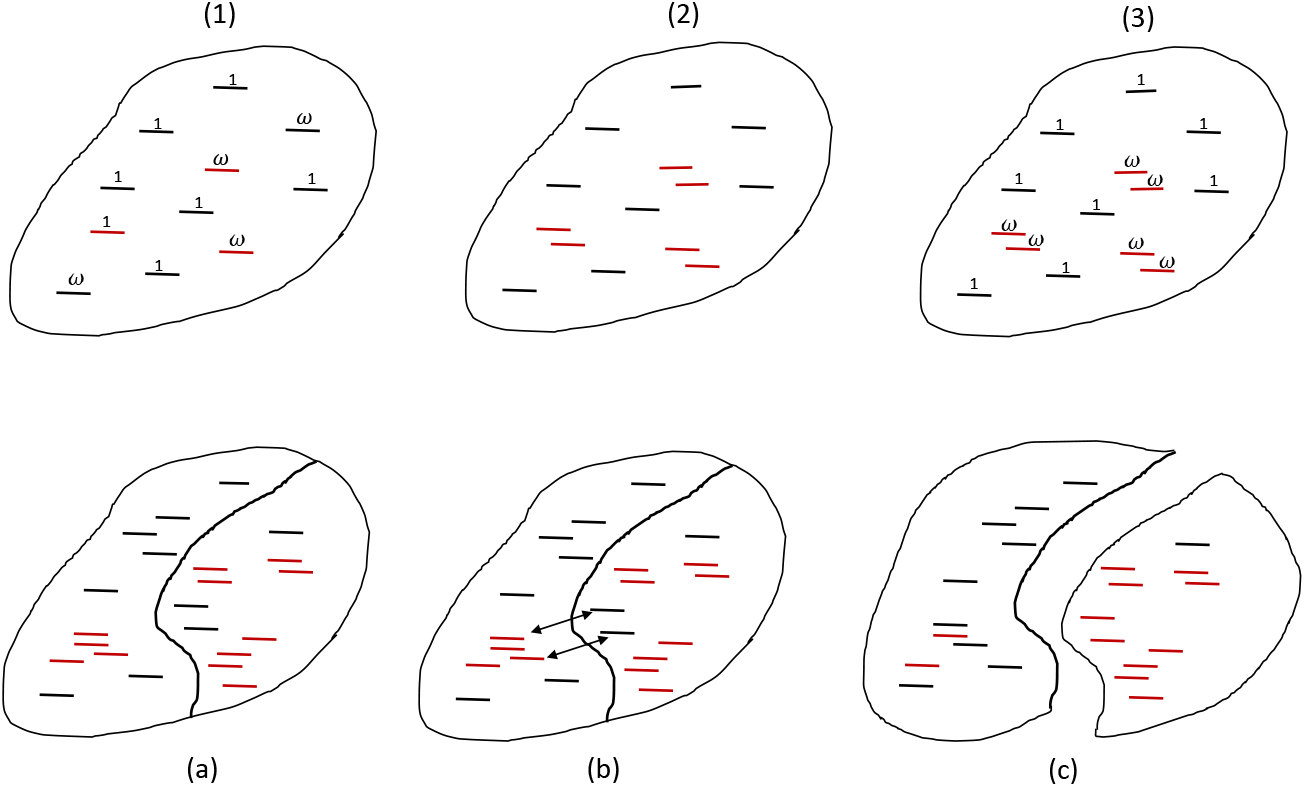
Steps of chromosomal replication (1-3) and the split (a-c) model. (1) selection of a set of *κ* chromosomes (in red) according to their dominance value *ω*, (2) replication of selected chromosomes and (3) reassign new dominance levels. (a) random split of chromosomes into two groups of equal size, (b) exchange *s* chromosomes between the two groups to increase the number of dominant mt in one of the two groups, and (c) fission of the cell into two, each carrying one group of the chromosomes.

By varying the values of these parameters, the model encompasses a wide range of mechanisms that potentially govern the binary fission of ciliates or a subset of ciliates, such as Tetrahymena. For example, the case with *κ* = 45 (assuming a 45-ploid macronucleus) corresponds to the model that all chromosomes in a macronucleus are replicated exactly once before the split. This is the common feature for prokaryotic binary fission and presumably also true for some eukaryotic cells. In such a situation, the dominance levels associated with chromosomes in the model have no effect. Also notice that regardless of the value of *κ*, there is no head-start effect when *ω* = 1 since all existing chromosomes have an equal probability of being selected. Aside from the extreme cases, various degrees of head-start mechanisms^30,33,39^ can be achieved by some combinations of *κ* and *ω*. In general, the smaller the *κ* and larger the *ω*, the stronger the head-start becomes. An example is *κ* = 1 and *ω* ≫ 1. In such a setting, a single chromosome which is randomly selected initially and its replicates will be the sole templates for all replications (45 rounds). Head-start increases the level of variation in the final outcome and its strength can be summarized by the probability that the next batch of replications will use exclusively the chromosomes and their replicates from the immediate previous batch. For example, with *κ* = 1 and *ω* = 100, the chromosome in the current batch and its replicate have probability 2 × 100/(2 × 100 + *k* − 2) of being the one used in the next round of replication, where *k* is the number of chromosomes after the current round of replication. After the first batch of replication, *k* = 46 and the probability is 0.82. Immediately before the last batch of replications, *k* = 89 and the probability becomes 0.70. Table 1 lists a few models of interest.

**Table 1.** Contrasting models of macronucleus proliferation in Tetrahymena

| Specific Models | Parameters | Key feature |
| --- | --- | --- |
| a: Null model | $\kappa = 45, \phi = 0$ | Even replication & random split |
| b: Random-split | $\phi = 0$ | Fission is random |
| c: No-head-start | $\omega = 1$ | No head-start effect |
| d: Extreme head-start | $\kappa \leq 2$ | Head start with a small $\kappa$ value |

Unlike the other two factors that may change the proportions of mts in a clonal population, deviation from random fission of a fully grown set of chromosomes in a macronucleus only has the impact of speeding the pace of purity by pushing more chromosomes of the same mt into the same daughter cells. One can go even further by allowing different parameters at different stages of development. For example, it may be postulated that the mechanism governing the chromosomal replications before the first cell division and after may be different.

## The experiment and observations

The experiment reported in this paper was designed to track the dynamics of mating type distribution of a clonal population founded by a single karyonide since such outcomes reflect the inner working of the binary fission. Each replicate of the experiment started with a single karyonide, and its asexual growth was monitored. At each of the pre-defined number of cell divisions, a sample of cells was taken from the clonal population, each of which was then cultivated separately to maturity to determine if it is pure for some mts. It should be noted that similar experiments focusing on either very early or late stages of growth of a clonal population were extensively conducted decades ago^32,38–40^. A detailed diagram on the experimental procedure is given in the Methods section.

For each replicate experiment tracking the clonal population with a single karyonide, we intended to take a sample of size 24 cells at each of the three time points 8, 30 and 90 divisions. Each cell in a sample was cultivated separately to form a sub-clonal population until maturity and their mt status were determined. Due to the failure of some sub-clonal populations, the sample size was typically smaller than originally anticipated. To compensate for the loss, sample size was doubled and two additional sampling times (20 and 50 divisions) were added in the last batch of the experiment. At the end, a total of 4704 sub-clonal populations were cultivated and examined, resulting in 3883 successful completions. Table 2 gives the results of the experiments for the three sampling times that were shared by all replicates. Since mt I is absent in the strain used in the experiment, only *n*_2_, …, *n*_7_, *n*_8_ are given in Table 2.

**Table 2.** Occurrences of mating types (*n_i_, i* = 2, …, 8) in samples from three sampling times

| Clone | 8th division | 30th division | 90th division | #mts |
| --- | --- | --- | --- | --- |
| a2B1 | 0,0,0,2,0,2,18 | 0,12,0,0,0,9,0 | 0,10,0,2,0, 7,0 | 3 |
| a3A1 | 0,0,0,0,2,0,17 | 0,0,11,1,2,9,1 | 0,0,10,0,4,6,0 | 4 |
| a3A2 | 0,0,0,0,0,0,23 | 0,0,0,0,0,0,0 | 1,0,6,0,4,12,0 | 4 |
| a3B2 | 0,0,0,0,9,0,13 | 0,0,2,0,1,15,6 | 0,0,3,0,15,3,0 | 3 |
| a4A2 | 0,0,3,0,0,0,18 | 0,0,15,0,2,4,3 | 0,0,13,0,6,3,0 | 3 |
| a4B1 | 0,0,3,0,0,1,19 | 0,0, 15,0,6,0,1 | 0,0, 14,0,0,9,0 | 3 |
| a4B2 | 0,0,0,0,19,0,5 | 0,0,0,0,20,0,3 | 0,0,0,0,18,3,0 | 2 |
| b1A1 | 0,0,2,0,1,0,14 | 0,0,6,0,4,3,2 | 0,0, 9,0,0,3,0 | 3 |
| b1A2 | 0,0,1,1,0,0,8 | 0,0, 13,1,0,0,3 | 0,0, 8,1,0,3,0 | 3 |
| b2A1 | 0,0,0,0,0,0,18 | 0,0, 12,0,4,1,6 | 0,0, 6,0,4,0,0 | 3 |
| b2B1 | 0,0,0,0,0,0,15 | 0,0, 20,0,1,0,1 | 0,0, 12,0,0,0,0 | 2 |
| b3A1 | 21,0,0,0,0,0,0 | 0,0,10,0,0,0,1 | 12,0,0,0,0,0,0 | 2 |
| c30B1 | 0,0,0,0,7,7,28 | 0,0,18,1,14,8,2 | 0,0,4,0,29,14,0 | 4 |
| c30B2 | 0,3,7,0,0,3,32 | 0,0,25,0,1,13,0 | 0,0,32,0,0,14,0 | 4 |
| c38A1 | 0,4,6,4,4,4,19 | 0,0,26,2,0,13,0 | 0,0,34,12,0,0,0 | 5 |
| c38A2 | 0,0,19,0,0,5,12 | 0,0,32,2,0,5,0 | 0,0,38,0,0,9,0 | 5 |

Note: An additional 26 clonal populations were observed to be pure for a single mt, 13 of which were IV, 6 were II, 4 were VI, 2 were V and 1 was VII. Clonal population names started with a,b and c represent 1st, 2nd and 3rd batches of experiments, respectively. #mts is the number of distinct mts observed in the collection of samples (including those not presented for c30B1, c30B2, c38A1 and c38A2). The percentage of clonal population with 1,2,3,4 and 5 mts were 62,7,17,12 and 2, respectively.

## Likelihood of the model parameters

Suppose a clonal population starts with a single karyonide. The status of the macronuclei in a random sample of *n* cells taken after the t-th divisions can be represented by the row vector **n**(**t**) = (*n*_1_(*t*), …, *n*_8_(*t*)), where *n_i_*(*t*)(*i* < 8) is the number of cells pure for mt *i* and *n*_8_(*t*) is the number of non-pure cells which eventually sort to generate more than one type of pure cells. Similarly the composition of the population can be represented by a row vector **N**(**t**) = (*N*_1_(*t*), …, *N*_7_, *N*_8_(*t*)) where *N_i_*(*t*)(*i* < 8) is the number of cells that are pure for mt *i* and *N*_8_ is the number of non-pure cells. The probability of observing **n**(**t**) given the underlying population composition **N**(**t**) can be derived exactly or be approximated by a multinomial distribution (see Methods).

When observations are available from more than one replicate experiment, the joint probability of observing all the data **D** is the product of the probability for each experiment, which leads to the likelihood function

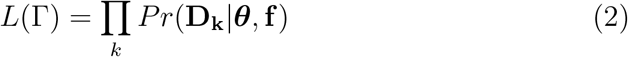

where **D**_**k**_ is the observation from the *k*th replicate,***θ*** is the vector of parameters defined by (1), **f** = {*f*_1_, …, *f*_7_} with *f_i_* being the probability that the initial macronucleus has *i* mt chromosomes, and the product is taken over all replicates. We set the initial state to be the composition of mt chromosomes of a macronucleus at octoploid. This likelihood function of a given ***θ*** and **f** can be evaluated numerically through Monte-Carlo simulation (see Methods).

## Inference on *θ* and f

Although the likelihood value for a given set of parameter values is not difficult to compute (i.e., estimate with sufficient precision), it is a challenge to search for the maximum likelihood estimate due to the very large number of combinations of parameters. We adopted a stepwise search strategy in which a preliminary search with a modest number of repeated simulations was conducted to identify promising local regions of parameters, and then gradually increase the simulation replicates to improve accuracy. It was found that during the first step the promising ranges of parameters were confined by *ϕ* ≤ 6 and *κ* < 10.

Furthermore, since each simulation of a clonal population started with an initial state at octoploid, **f** needs to be factored in to derive the final likelihood value. Fortunately this optimization problem can still be solved by a brute-force approach, which examined a very large number of combinations of parameters. Once the promising combinations of parameters were identified, their likelihood values, as well as those in their close vicinity, were re-estimated with a very large number of replicate simulations (usually several or even tens of millions) to ensure sufficient accuracy.

It is found that within the whole parameter space the global maximum likelihood is achieved at

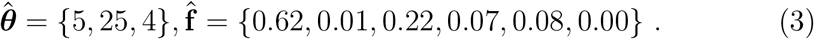

Since the peak surfaces for different values of *ϕ* hardly overlap for *κ* and *ω*, it allows one to present the information compactly. Figure 2 shows the log-likelihood surface within 1% of the maximum. One can see that there are three peaks of nearly the same height, followed by two lesser peaks.

**Figure 2.**
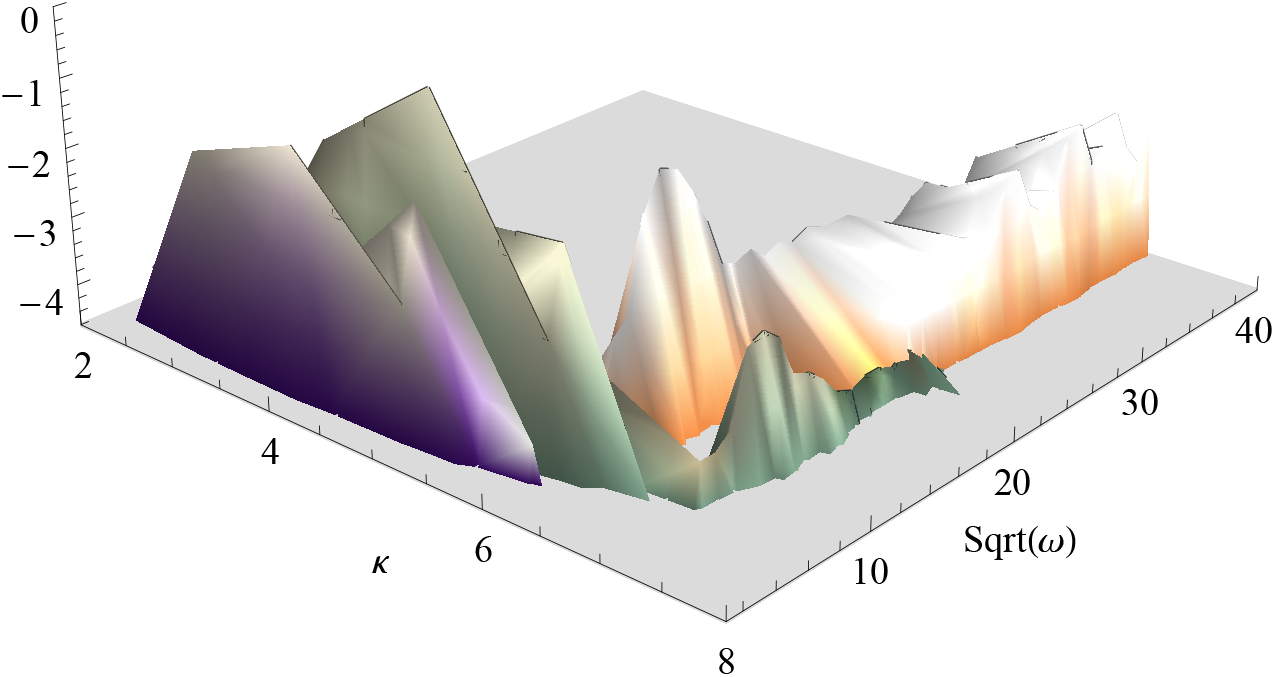
The combined likelihood surfaces within 1% of the maximum. Legend: orange for *ϕ* = 3, green for *ϕ* = 4, and purple for *ϕ* = 5. The highest peaks correspond to *ϕ* = 4, 3 and 5 are ***θ*** equal to {5, 25, 4}, {6, 144, 3}, {4, 9, 5}, respectively, with (5, 25, 4) being the highest.

The support for a number of hypotheses including the major peaks seen in Figure 2, judged by the log-likelihood ratio test, are shown in Table 3. It follows that the differences among the top three peaks in Figure 4 are not statistically significant. The null model (model a in Table 1) is so much inferior that it must be rejected outright. The hypothesis of random split (model b) regardless of the degrees of head-start is also soundly rejected, and mild deviation from the random split (*s*_1_ and *s*_2_) as well as severe bias (*s*_6_) can all be rejected, as is the no-head-start model rejected.

**Table 3.** Log-likelihood ratio tests and their significance for various models

| Model | Specific | $-2\ln(L/L_0)$ | Significance |
| --- | --- | --- | --- |
| 1 | $\theta = \{5, 25, 4\}$ | 0.0 | 1 |
| 2 | $\theta = \{6, 144, 3\}$ | 2.34 | $> 0.05$ |
| 3 | $\theta = \{4, 9, 5\}$ | 2.20 | $> 0.05$ |
| a | $\kappa = 45, \phi = 0$ | $> 1000$ | $< 10^{-20}$ |
| b | $\phi = 0$ | 61.0 | $< 10^{-15}$ |
| c | $\omega = 1$ | 60.5 | $< 10^{-15}$ |
| d | $\kappa \leq 2$ | 8.02 | $< 10^{-2}$ |
| $s_1$ | $\phi = 1$ | 30.7 | $< 10^{-8}$ |
| $s_2$ | $\phi = 2$ | 10.9 | $< 10^{-3}$ |
| $s_6$ | $\phi \geq 6$ | 10.5 | $< 10^{-3}$ |

Note: Models a,b,c and d were the same as those in Table 1.

To guard against unforseen failure of the models to capture the essence of the proliferation process, we compared the predictions of the top models with some features of the observations. One obvious characteristic is the percentage of purity at each sampling point among clonal populations that contain more than one mt.

Figure 3 shows the percentage of pure cells in the experiment and the cumulative probabilities of purity under several sets of parameters. The model corresponding to the optimal parameter clearly agrees with the observation.

**Figure 3.**
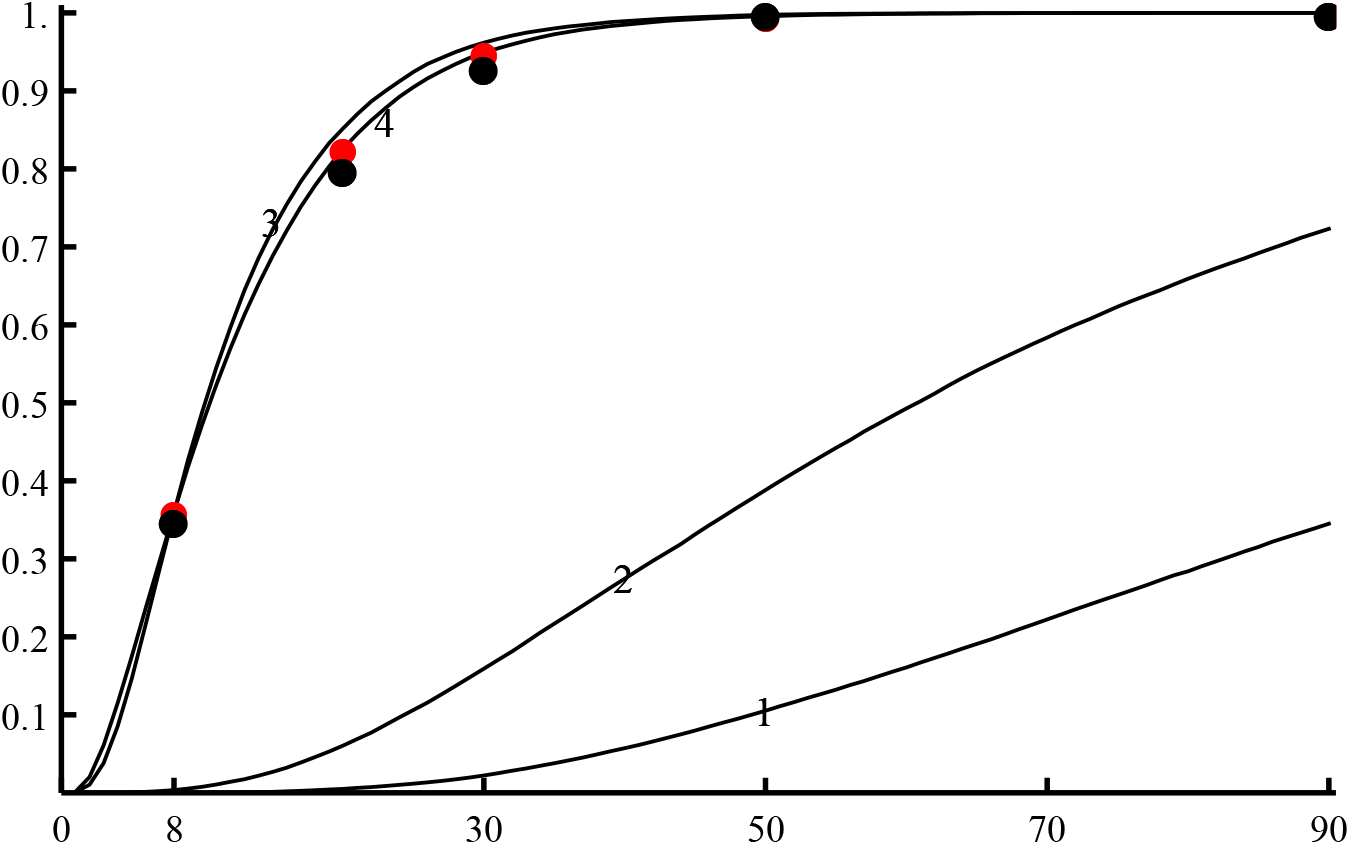
Comparison of the cumulative probabilities of purity under several sets of parameters and the observed proportions of purity. Legend: black dots: observed proportion, 1: ***θ*** = {45, 1, 0}, 2: ***θ*** = {4, 1, 0}, 3: ***θ*** = {4, 400, 0}, and 4: ***θ*** = {5, 25, 4}(line with red dots).

Our analysis shows that there is overwhelming evidence of a fairly strong head-start mechanism which gives numerical support to the head-start hypothesis first proposed by Orias and Baum^39^ which was put forward primarily for interpretation of the high proportion of purity at the time of maturity. That best models occur at *κ* = 5 is intriguing since there happens to be five chromosomes in the micronucleus. It is not clear if this is a coincidence or if the organism is somehow optimized to have the number of current replication processes roughly equal to the number of chromosomes in the micronucleus. In our formulation, the head-start mechanism has two parameters, one is the concurrent duplication process and the second is the dominance assigned to newly used templates and their product. Around the optimum, the dominance are all relative high. With *κ* = 5, *ω* = 25, the 10 chromosomes (5 templates and 5 replications) will collectively have dominance equal to 10 × 25 while the remaining 40 chromosomes only have a sum of dominance values equal to 40. Therefore, in the next round of replications, previously used templates and their replicates will have a large chance to be used again (within the 10 chromosomes, selection is equal). To be more precise, the probability that the next cycle of replications uses exclusively the 10 chromosomes has a probability equal to 0.48. A relative small number of concurrent replication apparatuses appear to make sense since it allows cells not to commit too many resources to the process of replications of many mini-chromosomes simultaneously. Such a strategy may be particularly important for some ciliates with extremely high ploidy copy^41^. By all means, models with *κ* > 8 have little support from the data. On the other hand, too few (*κ* ≤ 2) concurrent replication apparatuses do not agree with the data either.

That the likelihood of an observed pattern with a random-split is at least 10^15^ times smaller than that by the model with a non-random split leads to strong support for the non-random fission process. Therefore the random split assumption should be abandoned completely. Furthermore the degree of non-randomness is considerable since the case with *ϕ* = 1 will be at least 10^5^ times less likely than the optimum which occurs at *ϕ* = 3. The inability of a random split to generate the observed degree of purity is also seen by an analytical analysis (see Methods). This finding is in sharp contrast to the earlier analysis^26–28^ which concluded that fission was at random (in particular for the chromosome containing a heat sensitive gene). We suspect that in addition to a potential difference for a different genomic region, the high resolution of our data, the sophistication of our model and rigorous inference allowing the head-start of chromosomal duplication, contributes to the difference. Our conclusion follows as a logical consequence once it is accepted that there is a strong preference for continuing the replication process using the same templates or the newly arisen chromosomes, which suggest the importance of the proximity of chromosomes. It is reasonable to assume that chromosomes that are derived from the same founding ones (thus of the same mt) will have a tendency of aggregating together. When a cell splits into two, chromosomes that are physically closer to each other will thus have a higher chance of being pushed into the same daughter cells. However, despite the fact that evidence of a non-random split is strong, it also disfavors strongly a highly biased splitting.

By introducing the initial state of mts composition at the octoploid into the inference framework, we were able to separate the issue of mt sorting during the late stage of macronucleus development and binary fission from the issue of formation of mts in the early stage of macronucleus development. We found that the frequency of the numbers of mts does not impact the relative pattern of likelihood values of other parameters in the vicinity of the optimum, but has a significant impact on the maximum of the likelihood surface. On the other hand, our maximum likelihood estimate of the initial frequencies of mts provides a useful starting point to explore the mechanisms of their generations. The most striking features that need to be reconciled is the high frequency of purity even at the octoploid stage. The observation of more than 50% single mt clones were observed in many classical studies^38,42^, but these studies were focused on the composition of mts of a population at the stage of maturity which happens usually after 60 or more cell divisions. Therefore, those classic results were thought to be partially due to the fact that a sizeable portion of such cases were due to the sorting process. Given the likely high frequency of a single mt at octoploid, there is a need to explain how such a high frequency of a single mt at octoploid can arise. Alternative explanations were put forward before, including the “intranuclear coordination”^30,38^. The assumption that the sorting process starts from octoploid onward is not an intrinsic requirement of our analysis and can be pushed earlier or later.

In our analysis, we focused on the mechanism of binary fission and sorting of mts, we did so by disregarding difference in frequencies of different mts initially. This is equivalent to assuming that once created, the sorting of mts in a clonal population only depends on the number of mts and their relative frequencies at octoploid, but independent of detailed mts. Since the mt locus is silent during the asexual growth, this assumption should be reasonable. When ample data are available, the validity of this assumption can be evaluated. The final caveat is that our inference applies only strictly to the mtd chromosome in the macronucleus. Therefore, to what extent our conclusions about the binary fission of *T. thermophila* apply to other chromosomes as well as other Tetrahymena species or more distant ciliates remains to be illuminated.

As was pointed out earlier, it is conceivable that the mechanism of chromosomal replications before and after the first cell division may be different because the former starts with a diploid while the latter starts with 45 ploids. We performed additional analysis allowing for the possibility but did not find any evidence that this flexibility provides a better fit to the data. In particular, we explored the scenario in which the early chromosomal replications is governed by a stronger head-start (smaller *κ* and larger *ω*) coupled with a less biased split or random split which was inspired by earlier work^28^, and found that such models do not have sufficient support from our experimental data.

## Conclusions

Our primary conclusions is that the proliferation of the macronucleus of *T. thermophila*, for the locus of mating type, is governed by a strong head-start process in the replications of chromosomes, and considerable degrees of nonrandom split in the fission of the fully grown macronucleus. More specifically, the head-start has about five chromosomes being replicated at a time and a strong tendency to reuse the chromosomes involved in the early replications, and that the fission of a fully grown macronucleus is far from random, with a strong tendency to push chromosomes and their replications to the same daughter cell. Models representing either the conventional view extended from prokaryotic binary fission or synthesis from decades of classical experiments on *T. thermophila* are rejected.

Although these conclusions have overwhelming support from the data, it should be pointed out that different macro-chromosomes of *T. thermophila* may be subjected to different strategies due to various constraints, including varying size and order of replications, and further delineation is necessary to provide a comprehensive understanding of the proliferation strategies of *T. thermophila*. It may not be far reaching to suggest that other Tetrahymena species (or even in general ciliates) may employ similar/comparable strategies of cell proliferation. The illumination of the similarity and diversity of proliferation strategy across ciliates will be highly informative as it may have ramifications on the process for high polyploidy cells in multicellular organisms,including some human cancer cells, that acquire high ploidy due to endoreplication as briefly described in the beginning of the paper. A caveat of this study is the assumption that amitosis leads to equal copy number (i.e., 45) of each chromosome being passed to both daughter cells, which may not be entirely true^43^. Although we do not anticipate qualitative difference in major conclusions for a modest deviation from the current assumption, more elaborate analysis incorporating the uncertainty in chromosomal copy number can be performed when there exist sufficient experimental observations for specifying a reasonable model.

## Acknowledgements

The work was supported by the National Natural Science Foundation of China (no. 9163130009 YXF and no. 91231120 WM). We thank E. Orias for stimulating discussions and critical comments, and Sara Barton for editorial assistance.

## Author contributions

The project design: YXF and WM; experiment design: GYW, WM and YXF; experiment: GYW,KC and XFM. Modeling, inference and draft of the manuscript: YXF; refinement of inference and writing: YXF, WM,GYW and SQL.

## Methods

### The experiment design

The parental strains used for the experiment, expressing mating type IV and VI respectively, were generated through two rounds of genomic exclusion crosses between *T. thermophila* strains SB210 and B* VII (obtained from the Tetrahymena Stock Center at Cornell University). To induce conjugation, 15 ml equally mixed parental cells at a density of about 2 × 10^5^ cells/ml were starved in 10 mM Tris-HCl (pH 7.5) buffer in 250 ml Erlenmeyer flasks at 30°*C*. Individual conjugating pairs were isolated at the 14h post-mixing. When two exconjugants (named A and B) from a single pair separated, each was placed in separate drops (45ul) of SPP medium and examined frequently. After the first fission of each exconjugant, four newly developing macronuclei from the same pair are distributed to four daughter cells (karyonides, named A1, A2, B1 and B2). Each karyonide was first isolated into a drop of SPP medium and propagated for about 8 fissions. After isolating a sample of single cells into 96-well plates from the initial clonal population, the remaining cells were then inoculated into 20 ml tubes containing 5 ml fresh SPP medium and underwent another 12 fissions. Then serial transfer was performed by daily transfer with 1 : 2^5^ dilution in tubes with fresh medium. Each clonal population started with a single karyonide was cultured for a total of 90 fissions at 30°*C*, with shaking at 135 rpm. A total of 42 replicates were completed in three separate batches. A sample of 24 cells were planned initially at each of the three sampling points, 8, 30 and 90 divisions, for each clonal line. Since some of the selected cells failed in subsequent cultures and it was found that some existing mts segregating in the population were not discovered, we decided to double the sample size and had two additional sampling points for the third group of experiments to minimize the chance of non-detection of existing low frequency mts segregated in the population. Details of the experimental procedure is illustrated in Figure 4.

**Figure 4.**
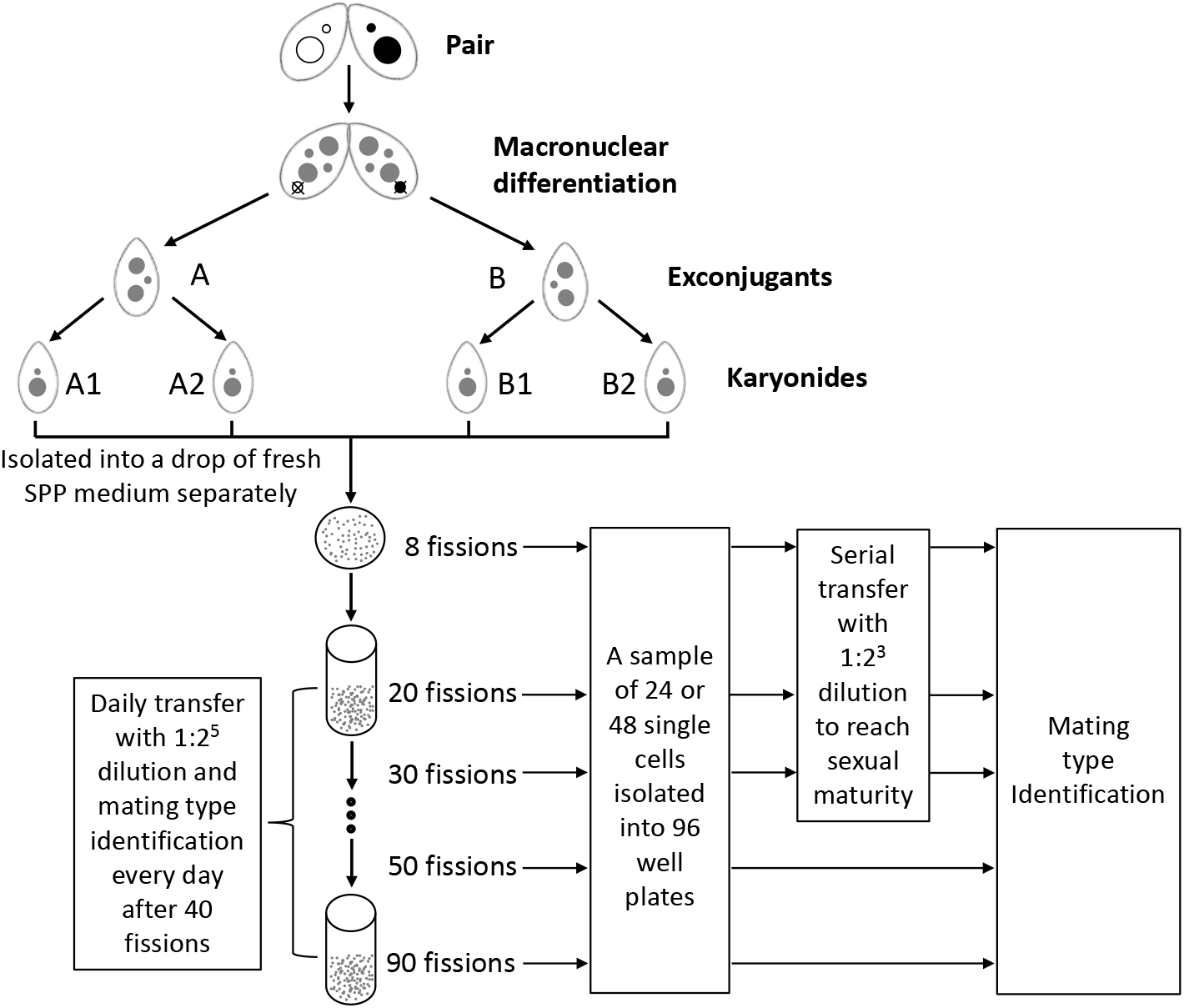
Design of the experiment.

For each cell in a sample, the result of successfully completing the experiment is either purity (pure for one of the six mts) or mixture of more than one mt. The steps are as follows. First, after the clonal population was propagated for 40 fissions, it was subjected to starvation every day to trigger mating, if mating is observed, the cell is non-pure, and when no mating is found, the same process is applied to the mixture of lines taken at each sampling point. Only when none of these tests resulted in mating being observed, the clonal population initiated from a single karyonide is declared to be pure for one mt. This procedure gives rise to the declaration of purity with high confidence. Therefore purity status for a clonal population is considered as observations without error. In case of non-purity, the mts of single cells isolated at each sampling point were further identified via mating type test experiments^44^ to determine all possible mts in the clonal population.

### Maximum likelihood inference of the models

Suppose a clonal population starts with a single karyonide with *k* mts. The status of the macronuclei in a random sample of *n* cells taken after the t-th divisions can be represented by the row vector **n**(**t**) = (*n*_1_(*t*), …, *n*_8_(*t*)), where *n_i_*(*t*)(*i* < 8) is the number of cells pure for mt *i* and *n*_8_(*t*) is the number of non-pure cells which eventually sort to generate more than one type of pure cells. Similarly the composition of the population can be represented by a row vector **N**(**t**) = (*N*_1_(*t*), …, *N*_7_, *N*_8_(*t*)) where *N_i_*(*t*)(*i* < 8) is the number of cells that are pure for mt *i* and *N*_8_ is the number of non-pure cells. The probability of observing **n**(**t**) given the underlying population composition **N**(**t**) is represented by

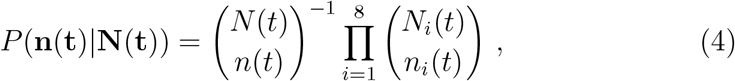

where *N* (*t*) is the total number of cells after the t-th division.

When there is no constraint, the population doubles in size every division, therefore when *t* is sufficiently large the population size will be large relative to the sample size. In such cases, the above probability can be well approximated by the multinomial distribution (with the convention that *a^b^* = 1 when *b* = 0, regardless of the value of *a*)

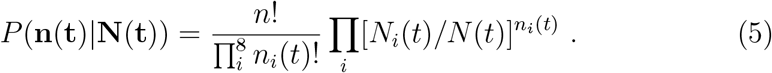

Although sampling in our experiment was without replacement, since the population size is sufficiently large (≥256) at the first sampling time while sample sizes (either 24 or 48) are relatively small, we will treat them as if they were taken with replacement. This treatment can lead to improved computational efficiency.

With multiple samples taken at times *t_i_, i* = 1…*s*, the joint probability conditional on the underlying population compositions is

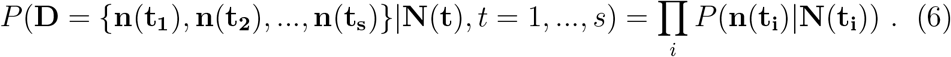

Since **N**(**t**) is not observed, one must consider all possible dynamics that can give rise to a particular combination of **N**(**t**), *t* = 1, …, *s*. That is equivalent to enumerating all possibilities of **N**(**t**), which leads to

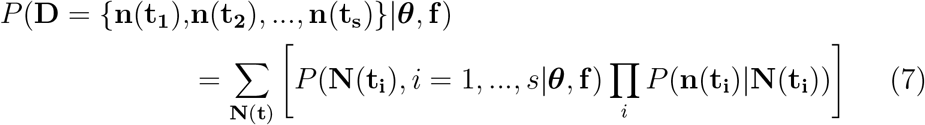

where ***θ*** is the vector of model parameters defined by (1) and **f** = {*f*_1_, …, *f*_7_} with *f_i_* being the probability that the initial macronucleus has *i* mt chromosomes. We set the initial state to be the composition of mt chromosomes of a macronucleus at octoploid. The rational is that given the observation in which up to five mts were segregated in a single clonal population, a considerable portion of the chromosomes in the macronucleus at diploid and tetraploid must not be committed, therefore octoploid is where all chromosomes can be committed in mt (although in reality it is still a small probability that full mt determination occurs later than octoploid).

When observations are available from more than one replicate experiment, the joint probability of observing all the data is the product of the probability for each experiment, which leads to the likelihood function

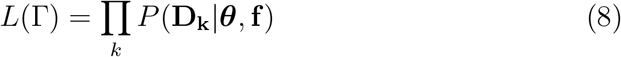

where **D**_**k**_ is the observation from the *k*th replicate and the product is taken over all replicates.

The probabilities (7)-(8) are untractable analytically and thus have to be evaluated only numerically. If one can simulate the propagation of a clonal population under a given model and an initial state, the probability (7) can be estimated by

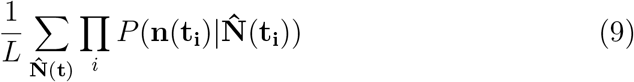

where 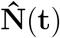 is a realization of **N**(*t*) in the total *L* simulations under parameters ***θ*** and **f**. The simulation process starts with the initial chromosome composition at the octoploid which grows to a full 45-ploid macronucleus. Each binary fission starts with replications of chromosomes from 45 copies to 90 copies, and then splits to obtain two daughter macronuclei. In more detail, once the value of ***θ*** is set, the binary fission process begins by randomly selecting a set of *κ* chromosomes, and then each is duplicated. The process is repeated until 90 chromosomes are obtained. However, after the initial round of chromosome duplication, the selection of a new set of *κ* chromosomes is governed by newly assigned dominance levels. For example, with *κ* = 1 and *ω* = 5, the original template and newly arisen chromosome will each have probability 2 × 25/(2 × 25 + 44) to be selected as the template for the next round of duplication. Once the macronucleus is fully grown, it starts the split.

This phase is simulated by first selecting a random split of 90 chromosomes into two halves, then increase the number of the most dominant mt between the two halves by up to *ϕ* by exchanging *s* randomly selected non-dominant mt chromosomes with *s* dominant mt chromosomes in the half. Since each round of binary fission results in doubling of population size, it is a challenge to keep track of the population. One effective way is to keep track of the entire population until the first sampling time *t*_1_ at which there are 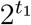 cells, then the composition of cells in a later sampling time is estimated through tracking a single descendant of a random subset of cells. This is done by first randomly selecting a cell out of the 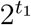 cells and following it through a given number of divisions, with only one daughter cell being retained after each division. Each realization of the history of a clonal population allows one to evaluate the probability of observed samples.

The precision of the above procedure for numerical estimation improves with the number of simulations. Any required accuracy can be achieved in principle by simulating a sufficiently large number of realizations. We found that with about half a million realizations, the numerical estimates become sufficiently stable, but we routinely simulated more than a million or more realizations to ensure adequate accuracy for any given set of parameters, particularly when these sets of parameters are close to the optimal solution. It should be pointed out that in addition to the parameters specified, the model is selectively neutral, in the sense that no mts or combination of mts have advantage over the other, except for the inherent difference in their frequency at the octoploid.

### Probability of purity under the null model

For a prokaryote with a single chromosome (mostly circular), the null model of binary fission is the only choice. Although this is also the common assumption for eukaryotic cells that undergo binary fission, its validity is not generally well scrutinized. For Tetrahymena, high ploidy makes the null model particularly challenging. To facilitate comparison, we explore one important property of this model.

Consider the mtd chromosomes in a *k*-ploid macronucleus. For simplicity, consider the situation of two mts present (without loss of generality type I and II) in the initial macronucleus. Suppose there are *m* copies of type I and *k − m* copies of type II. Let *q*(*l, i*) be the probability of having *i* copies of type I in a daughter macronucleus given there are *l* copies in the parent macronucleus.

Under the null model, the chromosomal replication phase results in two copies for each chromosome, and every split of the 2k chromosomes into two halves has the same probability which is 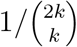. Therefore, the probability of a daughter macronucleus having *i* copy of mt I is

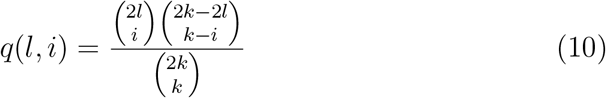

for 2*l* − *k* ≤ *i* ≤ *k* and 0 otherwise, where the nominator is the number of splits that can result in exactly *i* copies of type I chromosomes. It is easy to show that *q*(*l, i*) is symmetric for i around l and monotonically decreasing away from *l*. Let *q_t_*(*i*) be the probability that after *t* divisions, there are *i* copies of type I chromosomes. Then

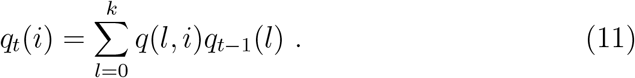

Purity is achieved when the number of type I chromosomes is equal to *k* or 0 (which means k copies of type II chromosomes). Therefore, the probability at division *t* is equal to *q_t_*(*k*) + *q_t_*(0). Figure 5 shows the probability of purity with regard to division *t* for several initial frequencies. It follows that when the proportion of the two alleles are about the same initially, the probability of purity is rather low after 30 divisions and only about 50% after 100 divisions. Relative high probability of purity at division 100 can only be achieved when the fixed mt was at extremely high frequency initially. This exploration illustrates that under the null model the probability of having high purity even at 100 divisions is very unlikely since the null model implies that initially the two mt chromosomes should have about equal frequency.

**Figure 5.**
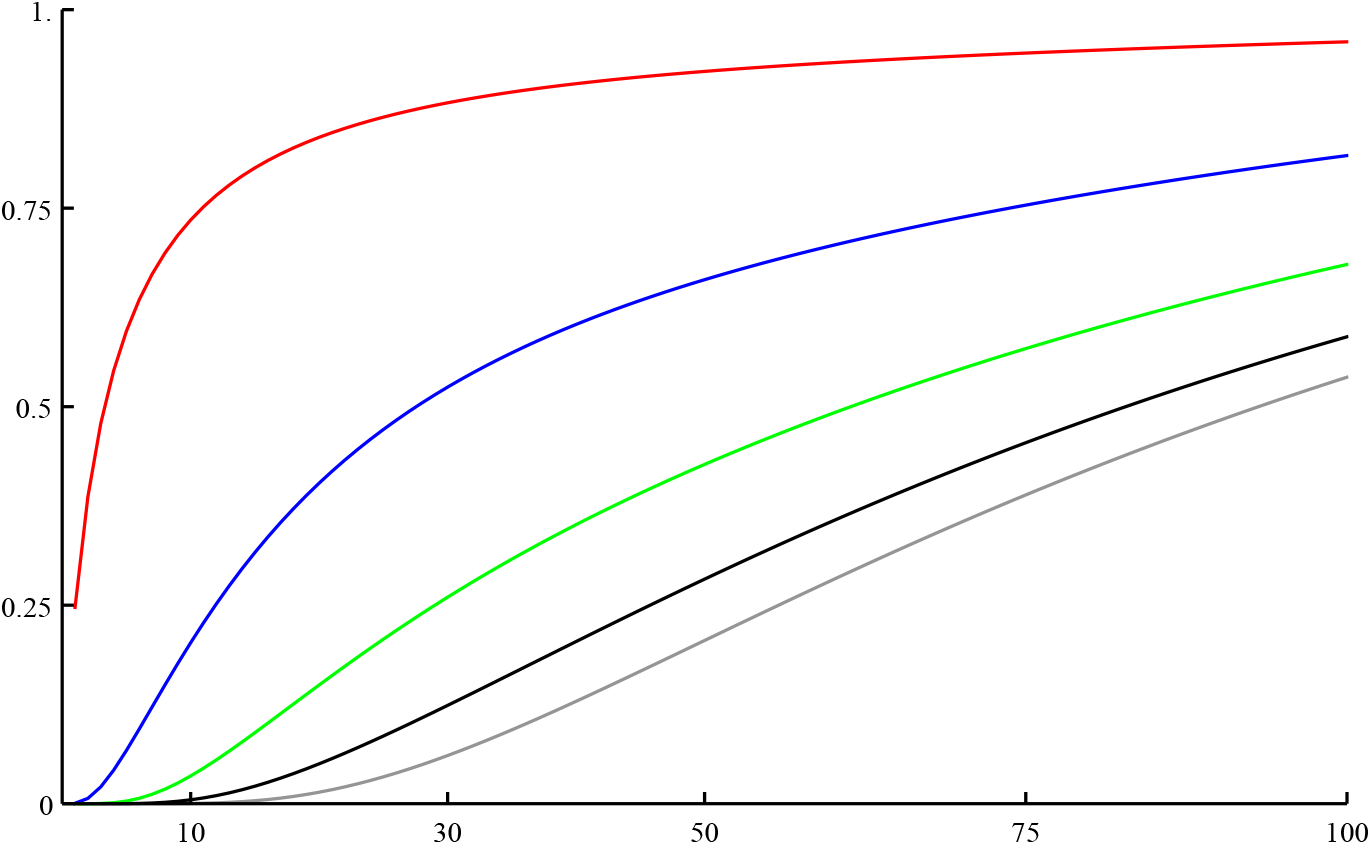
Probability of the purity of macronucleus at a given number of divisions with k=45. Lines in gray, black, green, blue and red correspond, respectively to the initial number of 23,30,35,40 and 44 alleles.

